# The first report on the effect of white and black truffle extracts on human semen parameters *in vitro*

**DOI:** 10.1101/2024.07.03.601846

**Authors:** Victoria Shelkovnikova, Maria Dmitrieva, Ekaterina Malygina, Natalia Imidoeva, Alexander Belyshenko, Maria Morgunova, Anfisa Vlasova, Tamara Telnova, Anna Batalova, Elena Martynova, Denis Axenov-Gribanov

## Abstract

Our experiment aimed to evaluate the influence of extracts of white and black truffle mushrooms on human spermatozoa. The study utilized 28 samples of wild truffle fruiting bodies. In the experiment, we used ejaculate from male volunteers of active reproductive age (N=10, 25–35 years old). During the experiment, we assessed sixteen physiological parameters. Research has shown that extracts from black and white truffles related to *Tuber* sp. have a stimulating effect on spermatozoa. The average path sperm velocity, curvilinear velocity and beat cross-frequency were increased by 56%, 48% and 50% respectively. Linearity showed a significant increase by 56% and straightness by 48%. This could be useful in the development of drugs to enhance sperm activity and lifespan. Additionally, extracts from black truffles have been found to have negative effects on spermatozoa, which could be relevant for developing new contraceptive drugs. Our study demonstrated, for the first time, the influence of methanol extracts of *Tuber* sp. mushrooms on male gametes *in vitro*.

## Introduction

Recently, truffle mushrooms have been considered not only as an exquisite food product but also as a source of natural compounds [1]. Truffle mushrooms are rich in biologically active molecules such as ascorbic acid, phenolic compounds, ergosterols, flavonoids, terpenoids, phytosterols, and polysaccharides. Due to this composition, truffles have the potential to be used as anticancer, antioxidant, antimicrobial, anti-inflammatory, and antidiabetic agents [2]. Recent studies have demonstrated that truffles have wound healing and antiviral properties [3]. Therefore, truffle mushrooms may have potential therapeutic applications.

Infertility is one of the most significant issues in modern healthcare. Approximately 13–20% of couples worldwide suffer from infertility, regardless of race or ethnicity. One of the main causes of male infertility is poor semen quality, such as low sperm concentration or motility [4, 5].

Surgical procedures and assisted reproductive technologies are employed to attain pregnancy in cases of infertility, but their drawbacks include high costs and limited accessibility for the general population [6, 7, 8]. Moreover, in some countries, there are no pharmaceuticals available to restore reproductive potential [9]. Medication therapy is limited to the intake of dietary supplements [10], which does not involve treatment for pathological conditions. Therefore, there is an urgent need for development of safe and effective means to improve semen quality both *in vitro* and *in vivo*.

Previous studies show that one of the components of truffle mushrooms is androsthenol related to the steroid group. This molecule increases in sexual desire [11]. Alcoholic extract of desert truffle *Terfezia boudieri* is known to significantly increase the levels of luteinizing hormone and testosterone in rats and, consequently, act as an aphrodisiac [12]. However, in 2014, it was found that when consuming truffle mushrooms as an aphrodisiac, attention should be paid to how truffles are prepared or cooked, their quantity, and stage of ripeness, as they contain flavonoids in the form of glycosides, which act as antagonists to male sex hormones [13]. Also, it is known that truffles often contain biogenic amines (serotonin, dopamine, and histamine) [14].

Another study demonstrated that hydroalcoholic extract of *Terfezia boudieri* exert influence on sperm and testosterone levels in male Wistar rats. The study revealed that rats injected with truffle extract experienced meaningful improvements in testosterone levels, body weight, testis weight, sperm count and sperm motility compared to the control group. These results suggest that *Terfezia boudieri* may play a significant role in addressing sexual impotence and infertility in males [15].

Truffle extracts possess unique properties that make them potentially valuable for medicinal purposes. The use of truffles in reproductive medicine may be of particular interest for development of medications with both spermatozoa-activating and contraceptive effects. Therefore, our study aimed to investigate the effects of truffle extracts on human spermatozoa in vitro.

## Methods

### 1. Collection of truffle mushrooms and preparation of samples

The study used 28 samples of wild truffle fruiting bodies of black (*Tuber macrosporum*) and white (*Tuber magnatum*) truffles collected in Krasnodar Region from August to November 2020–2022. The mushrooms were collected near the cities of Krasnodar, Sochi and Maykop (southern part of the Russian Federation). To prevent decay, the mushrooms were placed in plastic containers with soil and rice. During transportation, the samples were kept at 6–10°C. The received truffle mushrooms were cleaned with water and ethanol with a toothbrush. The mushrooms were then dried at room temperature for 5 hours to remove excess moisture, and stored at –20°C. Each truffle extract was assigned a unique identification number as presented in Table 1.

**Table 1.** Numbering of extracts obtained from truffle mushrooms.

| Collection location | Numbers of truffle mushroom extracts |
| --- | --- |
| Krasnodar | Black truffles: *T. macrosporum* LPB2020-16, *T. macrosporum* LPB2020-17, *T. macrosporum* LPB2021-26, *T. macrosporum* LPB2021-27, *T. macrosporum* LPB2021-28, *T. macrosporum* LPB2021-29, *T. macrosporum* LPB2021-30, *T. macrosporum* LPB2022-39, *T. macrosporum* LPB2022-40, *T. macrosporum* LPB2022-41, *T. macrosporum* LPB2022-42, *T. macrosporum* LPB2022-44, *T. macrosporum* LPB2022-47, *T. macrosporum* LPB2022-67, *T. macrosporum* LPB2022-68, *T. macrosporum* LPB2022-69, *T. macrosporum* LPB2022-70, *T. macrosporum* LPB2022-71, *T. macrosporum* LPB2022-72;<br>White truffles: *T. magnatum* LPB2021-31, *T. magnatum* LPB2021-32 |
| Sochi | Black truffles: *T. macrosporum* LPB2021-22, *T. macrosporum* LPB2021-23, *T. macrosporum* LPB2021-24 |
| Maykop | Black truffles: *T. macrosporum* LPB2022-58, *T. macrosporum* LPB2022-59, *T. macrosporum* LPB2022-61, *T. macrosporum* LPB2022-63 |

### 2. Extraction of biologically active components of truffles

For extraction, 0.5 g of each truffle sample was ground into powder using a mortar and pestle, with methanol (Vecton, Russia) added in a ratio of 1:10. The obtained mixture was incubated for one hour on a roller shaker MX-T6-S (BIOBASE, China, Jinan), and then centrifuged with LC-04A centrifuge (Armed, Russia) for 10 minutes at 3 000 rpm. The supernatant was transferred to different tubes [16].

### 3. Evaluation of the effect of natural compounds of truffle extracts on spermatozoa *in vitro*

In the experiment, we used ejaculate from male volunteers of active reproductive age (N=10, 25–35 years old). After being briefed on the research methods and objectives, each male participant provided a written consent for the study. The ejaculate samples were collected, by masturbation, in sterile containers after a period of abstinence for 3–5 days. The effect of truffle extracts on human spermatozoa was assessed using a microscope with a temperature-controlled stage PLS-MY-B041A-3 (PLSUP, China), “Semen and Sperm Quality Analysis System” software (V1.12), and a 96-well plate with the optically clear bottom. The analysis was performed in automatic mode. Physiological parameters of spermatozoa were measured according to protocols described in the 5th edition of the WHO Laboratory Manual for the Examination and Processing of Human Semen [17].

The experiments were conducted at 37°C. For control conditions, 25 µL of methanol was applied to a sterile 96-well plate and dried until completely evaporated. Subsequently, 150 µL of ejaculate was added to the evaporated methanol fraction.

The experimental samples were divided into 2 groups. For the first group, concentrated truffle extracts were used in amount of 25 µL. For the second group, the concentrated extract was diluted in a ratio of 1:6 (2.5 µL of extract + 12.5 µL of methanol). The extracts were dried until complete evaporation of methanol (solvent), and then 150 µL of ejaculate was added. The analysis was performed in three analytical replicates.

To prepare wet mount, we applied 10 µL of ejaculate onto a microscope slide and covered it with a 22×22 mm cover slip [17, 18]. The microscope stage was preheated to 37°C. During the experiment, we assessed sixteen physiological parameters, specifically: total of motile sperm, percent of motile sperm, fast progressive sperm (WHO A), slow progressive sperm (WHO B) local motile (WHO C), immotile (WHO D), average path velocity (VAP), curvilinear velocity (VCL), straight-line velocity (VSL), amplitude of lateral head displacement (ALH), beat cross frequency (BCF), percent of line moving, linearity (LIN), straightness (STR), wobble (WOB), mean move angle (MAD). Physiological parameters of spermatozoa were measured after 1, 3, and 6 hours of incubation at 200x magnification.

### 4. Statistical data processing

For statistical processing we employed the Past software (V4.03) using the ANOVA [19] analysis of variance with the Mann-Whitney criterion [20]. Differences between the mean values of the parameters were considered significant at p ≤ 0.05 [21].

## Results

The experiments revealed that after 3-hour incubation, no motile spermatozoa were observed in concentrated extracts. Of the diluted extracts, eight are noteworthy for their significant effects on reproductive cells. After 3 hours of incubation under the influence of diluted extract of the white truffle *T. magnatum* LPB2021-31, we observed a significant 56% increase in VAP (from 9.4±2 to 14.7±2.9 um/s). VCL and BCF also increased by 48% (from 14.5±3.4 to 21.5±3.7 um/s) and 50% (from 1.8±0.6 to 2.7±0.7 Hz) respectively (Fig. 1). In diluted extract of the black truffle *T. macrosporum* LPB2022-67 after 6-hour incubation, LIN showed a significant increase by 56% (from 40.7±11.1 to 63.6±14.8) and STR by 48% (from 58.3±8.8 to 86.5±20.1).

**Fig. 1.**
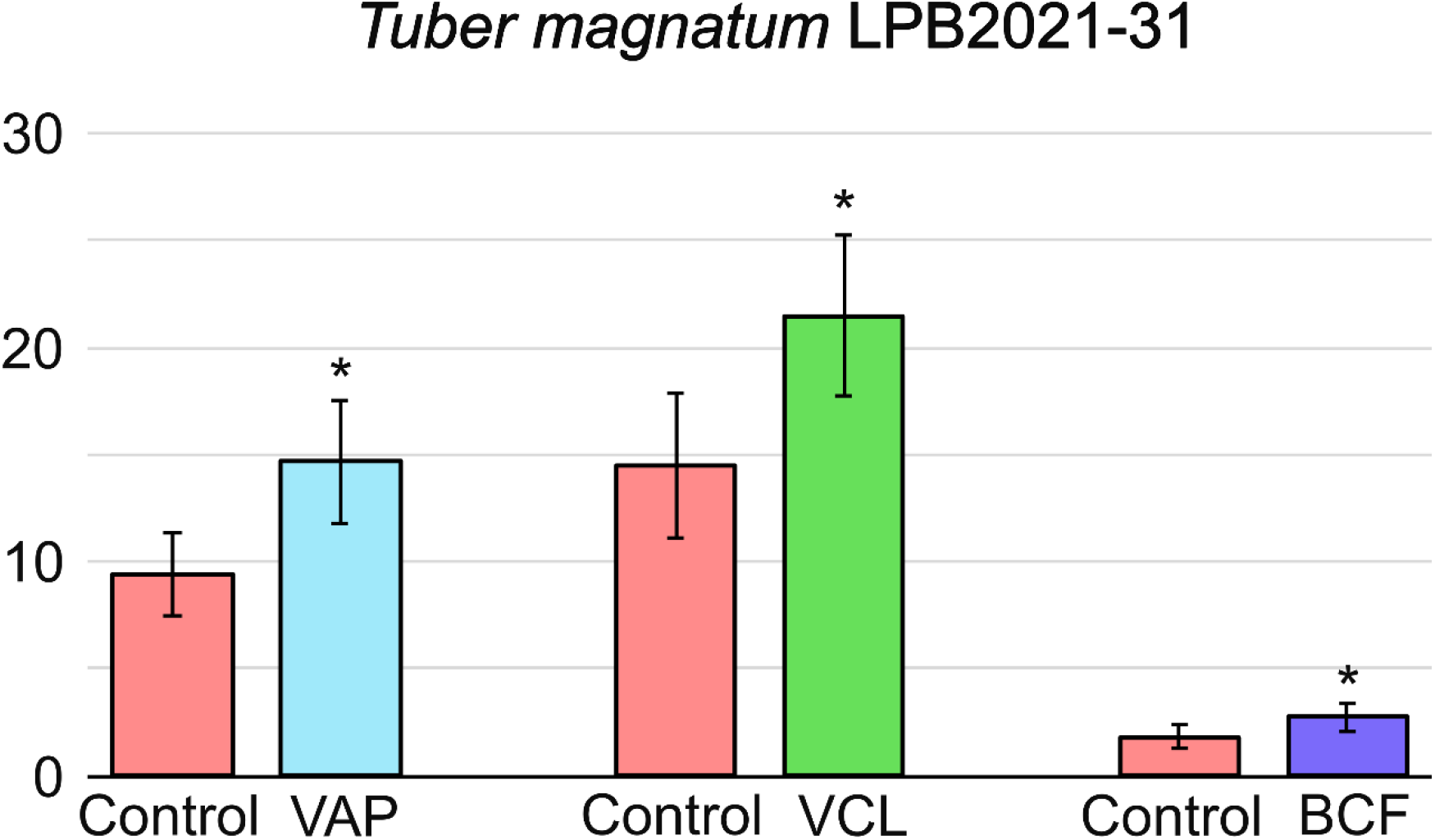
Histogram of extract of the white truffle T. magnatum LPB2021-31 stimulatory effects on spermatozoa in vitro. «VAP» is the average path velocity; «VCL» - curvilinear velocity; «BCF» - beat cross-frequency.

Certain truffle extracts had adverse effects on spermatozoa, resulting in reduction of their physiological parameters when compared to the control conditions. After 3 hours of incubation, we could see a significant decrease in the percentage of motile spermatozoa in extracts of black truffles: by 39% for *T. macrosporum* LPB2022-39 (from 96.2±7.7 to 58.9±3.1%), 33% for *T. macrosporum* LPB2022-61 (from 96.2±7.7 to 64.2±15.1%), by 24% for *T. macrosporum* LPB2022-63 (from 96.2±7.7 to 72.8±11.8%), and by 22% for *T. macrosporum* LPB2022-72 (from 96.2±7.7 to 75 ±8.3%). After 3 hours of the experiment, WHO C in black truffle extracts decreased considerably: by 51% for *T. macrosporum* LPB2022-61 (from 84.1±17.2 to 40.1±17.3 um/s) (Fig. 2) and by 41% for *T. macrosporum* LPB2022-69 (from 84.1±17.2 to 34.3±1.7 um/s). After 6 hours, STR in extract of black truffle *T. macrosporum* LPB2022-71 has also decreased by 52% (from 5.2±2.1 to 2.5±0.8 um/s).

**Fig. 2.**
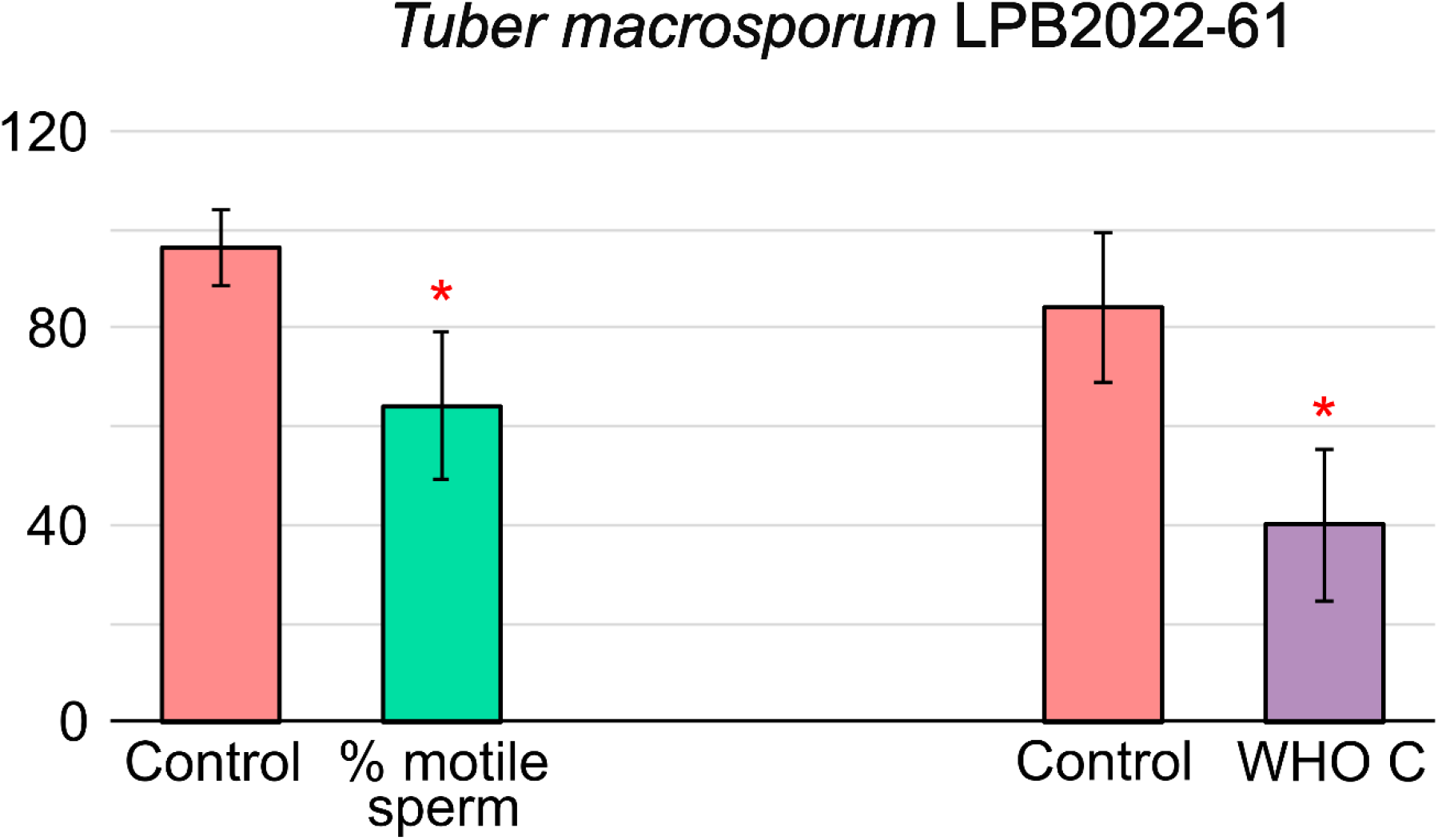
Histogram of negative effects of black truffle extract *T. macrosporum* LPB2022-61 on spermatozoa *in vitro*. «% motile sperm» - percent of motile sperm; «WHO C» - local motile.

Thus, according to first results of our experiments, the stimulating effect on spermatozoa in vitro was produced by extracts of black truffle *T. macrosporum* LPB2022-67 and white truffle *T. magnatum* LPB2021-31. Extracts of black truffles *T. macrosporum* LPB2022-61, *T. macrosporum* LPB2022-63, *T. macrosporum* LPB2022-67, *T. macrosporum* LPB2022-69, *T. macrosporum* LPB2022-71, and *T. macrosporum* LPB2022-72 had a negative effect on male germ cells.

## Discussion

Infertility is a condition of male or female reproductive system characterized by the inability to attain pregnancy after regular unprotected sexual intercourse for 12 months or longer. Male infertility is positioned to be incurable. Treatment of male infertility involves boosting immunity and taking dietary supplements (a mixture of plant extracts). The Dictionary of Natural Products (DNP) database includes 55 natural compounds with spermatozoid-activating activity. The main producers of these compounds are microorganisms. The latter represent a promising source of unique natural compounds for treating different human diseases. Secondary bacterial metabolites possess a variety of therapeutic activities, including antimicrobial, antiviral, antitumor, etc. [22].

In this study, we have shown that extracts of white and black truffle *Tuber* sp. are primarily distinguished by their spermicidal properties. Percentage of motile spermatozoa, local motility and straight-line velocity were significantly low. This feature could be utilized to develop new contraceptives. However, two extracts from *Tuber* sp. showed spermatozoa-activating effects. VAP, VCL, BCF, LIN, and STR were remarkably high. These are key parameters describing sperm motion. VAP measures the average speed of sperm along a smoothed trajectory taken by the sperm [23]. This averaging helps to eliminate sharp changes in direction and variations in speed. VAP is used to analyze the overall motility, as well as their potential to reach the oocyte [24]. VCL reflects the average speed of sperm along the curved path it takes [25]. This speed measurement of the actual path followed by the sperm is an important indicator of sperm activity [23]. High VCL values often correlate with increased sperm fertilization ability, as it indicates high motility and the ability to overcome obstacles on the way to the oocyte. BCF describes the sperm’s tail beat frequency. It measures how often the sperm tail crosses the average line of its trajectory [26]. Low BCF may indicate motor defects in sperm, which in turn can affect their fertilizing capacity [27]. LIN assesses how straight the sperm moves [28]. It is calculated as the ratio of the average speed along straight trajectory (VSL) to VCL. High LIN indicates that the sperm is moving straight towards the oocyte [29]. STR also measures the straightness of sperm motion, but unlike LIN, STR is calculated as the VSL to VAP ratio. This parameter helps to assess how effectively the sperm follows its path. High STR means that the sperm is moving directly towards the oocyte, minimizing deviation from the trajectory. In procedures such as IVF and ICSI, these parameters are used in selecting the most motile and potentially viable sperm. Thus, these parameters play a major role in diagnosing infertility [29].

The fruiting bodies of truffle fungi contain numerous symbionts, secondary metabolites of which exhibit various biological activities. Earlier, we demonstrated that gleba and peridium of truffle fruiting bodies have different chemical composition [14]. Truffles have a unique composition of bacteria that have not been described in European representatives of *Tuber* sp. [30]. Symbiotic relationships between truffles and different microorganisms can induce production of new specialized metabolites [31]. The observed biological activity of truffles can presumably be related to the synthesis of secondary metabolites by the bacterial consortium found in these fungi. This could explain unstable activity of extracts from different fruiting bodies, and the limited use of truffle fungi in modern European medicine.

In addition, the analysis of the problem using patents and inventions revealed several interesting references to truffle effects in Chinese medicine. Patent №CN109045095A describes a formulation that stimulates spermatogenesis, improves sperm count and viability, and alleviates fatigue. The composition includes concentrated maca powder (*Lepidium* sp.), extract of male silkworm moth, *Epimedium* sp., truffle mushrooms, and raspberry [32]. Another patent №CN108740670A describes the creation of a beverage based on peptides that act as potent aphrodisiacs. One of the components of this beverage is black truffle peptide [33]. Also, patent №CN112755171A describes the creation of a composition to improve male sexual function. The composition comprises three main components, one of which enhances sperm motility and improves prostate function. This component includes soy protein powder, black truffle, zinc, taurine, and lycopene. Thirty healthy men aged 30–50 years consumed the above composition three times a day for a week. As a result, 83.3% of them experienced a significant improvement in sexual activity [34]. These patents provide formulations containing complex mixtures, including numerous ingredients from traditional Chinese medicine, making it difficult to study the true effects of truffle mushrooms on male reproductive system.

The invention presented in patent №US2013295140A1 is based on truffle mycelium extract, which stimulates testosterone production through olfaction. The smell of truffles stimulates testosterone production in humans by activating androgen receptors in brain. Increased testosterone levels lead to improvements in spermatogenesis and ejaculate quality [35, 36]. These patents have repeatedly referred to the stimulating effects of black truffle extracts in different mixtures.

Thus, this study represents the first assessment of the influence of methanolic extracts of black and white *Tuber* sp. on male germ cells in vitro. Although it is the first study in this area, truffle extracts showed significant changes in physiological parameters of human spermatozoa compared to the control group. The results provide a meaningful basis for isolating active compounds and identifying natural compounds responsible for activating the reproductive potential. Therefore, truffle mushrooms have significant medical potential as agents in the search for new biologically active compounds.

## Acknowledgments

The study was conducted with the support of the Russian Science Foundation (project 22-76-10036).

## Author contributions

Conceptualization: MED, VNS. Methodology: VNS. Software: AAV. Validation: MED, MMM, DVA-G. Formal analysis: EVM. Investigation: EVM. Resources: NAI, EIM. Data curation: AYB. Writing –original draft preparation: VNS. Writing – review and editing: MED. Visualization: TYT, AAB. Supervision: DVA-G. Project administration: DVA-G. Funding acquisition: DVA-G.

